# The polar localization hub protein PopZ restrains adaptor dependent ClpXP proteolysis in *Caulobacter crescentus*

**DOI:** 10.1101/301606

**Authors:** Kamal Kishore Joshi, Peter Chien

## Abstract

In *Caulobacter crescentus*, timely degradation of several proteins by the ClpXP protease is critical for proper cell cycle progression. During the cell cycle, the ClpXP protease, the substrate CtrA and many other proteins are localized to the stalked pole dependent on a polar interaction hub composed of PopZ protein oligomers. Prior work suggests that the localization of ClpXP, protease substrates, and cofactors is needed for recognition of substrates such as CtrA by ClpXP. Here, we formally test this hypothesis by examining the role of PopZ in ClpXP activity and find surprisingly that CtrA degradation is enhanced in cells lacking polar localization due to loss of PopZ. The ClpXP adaptor CpdR is required for this enhanced degradation of CtrA and other adaptor-dependent substrates, but adaptor-independent substrate degradation is not affected upon loss of PopZ. We find that overexpression of PopZ also leads to faster degradation of CtrA, but is likely due to nonphysiologically relevant recognition of CtrA by ClpXP alone as loss of CpdR does not affect this enhancement. Our main conclusion is that loss of PopZ, and therefore loss of polar localization, does not result in the loss of ClpXP regulated proteolysis, as would be predicted from a model which requires polar localization of ClpXP for its activation. Rather, our data point to a model where PopZ normally restrains ClpXP proteolysis by promoting the inactivation of the CpdR adaptor, likely through the phosphorylation activity of the CckA kinase.

## IMPORTANCE

Regulated proteolysis is critical for the cell cycle progression of bacteria such as *Caulobacter crescentus.* According to one model, this regulated proteolysis requires localization of the ClpXP protease at the stalked pole for its subsequent degradation of substrates such as CtrA. This study offers evidence that supports an alternative model to explain how localization might influence protein degradation. Using a delocalized *in vivo* system created by the deletion of a pole organizing protein PopZ, we show that activation of the ClpXP protease is independent of its polar localization. The data points to a role for PopZ in restraining ClpXP activity, likely by controlling the activity of upstream regulators of protease activity, such as CckA, though changes in its localization.

## INTRODUCTION

Proteolysis plays an important role in facilitating cell cycle progression and various developmental transitions in bacteria. Examples are: cell cycle progression in *Caulobacter crescentus* (*Caulobacter* hereafter) and cellular transition from vegetative to sporulation stage in *Bacillus subtilis* (1–4). The cell cycle of *Caulobacter* starts with a replication incompetent motile swarmer cell (G1-like phase). In response to developmental cues, the swarmer (SW) cell differentiates into a stalked (ST) cell. The ST cell is capable of replication and division to give birth to a new SW cell. Following replication and asymmetric division the mother ST cell immediately starts DNA replication and another round of cell division whereas the new born SW cell again has to differentiate into ST cell in order to continue its cell cycle (5, 6). To maintain such stringent control during SW-to-ST transition, levels of many proteins, including a protein called CtrA, change dramatically (3, 7).

CtrA is a transcriptional factor that controls transcription of ~95 genes and also functions as an inhibitor of DNA replication (8, 9). In SW cells, CtrA is phosphorylated by a membrane-bound bifunctional kinase CckA via the phosphotransferase ChpT (10–12). The phosphorylated CtrA binds tightly to DNA at chromosomal origin of replication (*oriC*) to block the initiation of replication (3, 8, 12). The same CckA-ChpT kinase pathway also phosphorylates the ClpX adaptor and response regulator CpdR in the SW cell which functionally inactivates it (12–15). During the SW-to-ST transition, the same CckA-ChpT cascade dephosphorylates both CtrA and CpdR via shuttling of the phosphoryl groups back to CckA (16, 17). The dephosphorylated CpdR then drives localization of ClpXP protease to the stalk pole (13). In parallel with this, other cofactors, PopA and RcdA, were shown to localize CtrA to the same stalk pole (18, 19). This convergent localization of ClpXP protease and the CtrA substrate at the stalk pole was postulated to increase the local concentration of the protease and the substrate leading to the degradation of CtrA (19, 20). The destruction of CtrA then allows the assembly of replication machinery at *oriC*, which then initiates the replication process.

A pole organizing protein, called PopZ (Polar Organizing Protein-Z), was identified in recent years. PopZ functions as a scaffold to recruit the accessory factors and the protease complex involved in CtrA degradation at the stalked pole (21). Besides serving as a scaffold, PopZ also anchors sister chromosomes at the stalked pole by directly binding to ParB, which is associated with *oriC* during chromosome segregation (22, 23). It also mediates polar localization of proteins involved in cellular signaling including both the transmembrane histidine kinase CckA and the DivJ protein through SpmX binding (23, 24). It was proposed that PopZ functions as a switch between chromosome tethering and protein scaffolding during SW-to-ST transition in *Caulobacter* to accommodate programmed asymmetry during cell cycle (21). Further dissection of PopZ protein revealed an N-terminal region, which is sufficient for binding all its partner proteins, and a C-terminal region for homo-oligomerization (25). Together, these protein localization studies supported a model where spatial compartmentalization of the protease ClpXP and the substrate CtrA allows removal of CtrA during SW-to-ST transition in *Caulobacter.*

Studies that were conducted recently suggested that spatial localization might not be critical for CtrA degradation. The factor RcdA was shown necessary for CtrA degradation as deletion of this factor did not support cell cycle changes in CtrA degradation (18). Structure guided RcdA mutants that were incompetent to localize to the stalk pole also failed to localize CtrA, but this delocalization did not perturb cell cycle-dependent degradation of CtrA (26). Localization of ClpXP to the stalk pole is not necessary for some of its activity as some ClpXP substrates, such as FtsZ, were still degraded in a SW cell where ClpXP was shown to be delocalized (27). Furthermore, *in vitro* reconstitution experiments using highly purified proteins supported an adaptor hierarchy model where CpdR, RcdA and PopA work in a coordinated fashion as adaptors to degrade many substrates including CtrA (4, 15).

In this study, we find that CtrA degradation is enhanced in PopZ lacking cells suggesting that PopZ restrains CtrA degradation. The ClpXP adaptor CpdR is required for this enhanced degradation as CtrA degradation was stabilized in a Δ*cpdR*Δ*popZ* strain. This degradation enhancement in Δ*popZ* cells also extends to additional CpdR and RcdA-dependent substrates, PdeA and TacA, respectively. These results indicate that ClpXP is activated at the CpdR adaptor level. However, ClpXP activity is not globally stimulated in Δ*popZ* cells as degradation of adaptor-independent substrates, such as a degron-appended GFP, is not affected. Overexpressing PopZ also leads to enhanced degradation of CtrA, which we propose reflects a nonphysiological recognition of CtrA by ClpXP alone, as loss of the CpdR adaptor does not affect this enhancement. Together, these results support a model where PopZ-mediated localization of the protease ClpXP and its adaptors is not essential for its activation but rather PopZ may affect protein degradation principally through regulation of upstream indirect regulators of the ClpXP protease such as the CckA kinase.

## MATERIALS AND METHODS

### Bacterial culture conditions and plasmid construction

*E.coli* and *Caulobacter* strains used in the study are tabulated in Table 1. *E*. *coli* strains were grown in LB media at 37 °C with the appropriate antibiotic (50 μg/ml kanamycin, 15 μg/ml tetracycline, 30 μg/ml chloramphenicol). *Caulobacter* strains were grown in Peptone-Yeast-Extract (PYE) media at 30 °C with the appropriate antibiotic (1 μg/ml tetracycline, 1 μg/ml chloramphenicol, 5 μg/ml kanamycin). PdeA and PopZ was PCR amplified and cloned into pENTR plasmids. The insert was then moved into xylose-inducible expression plasmids which append an M2-epitope tag on the N-terminus of the protein using Gateway-based cloning (28). Microscopy was performed as described previously (4, 15).

**Table 1.**
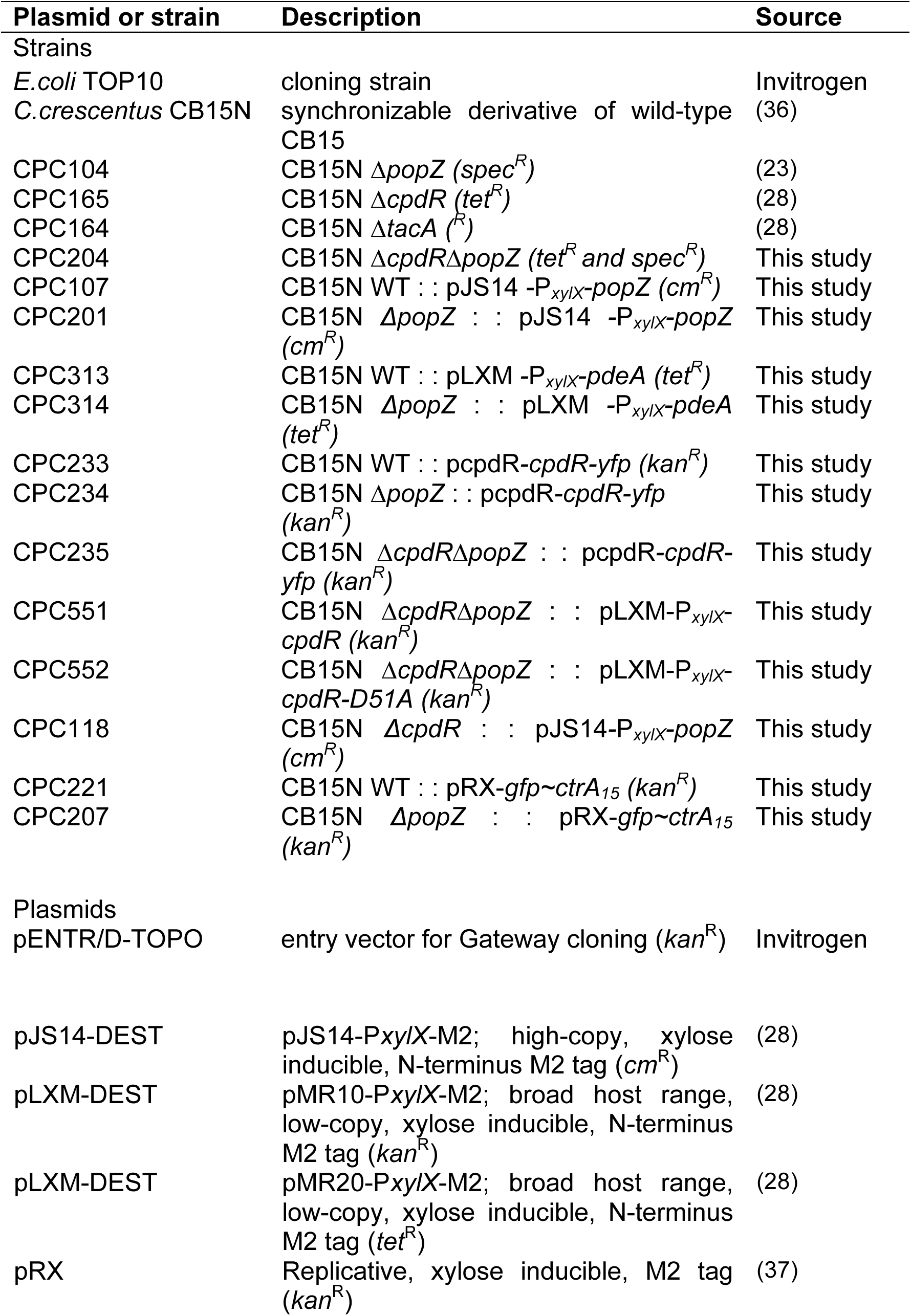
Strains and plasmids used in this study.

### *In vivo* protein degradation assays

*Caulobacter* cells were grown to OD_600_ ~ 0.3 in PYE medium with appropriate antibiotic. Protein expression was induced with xylose wherever required and then protein synthesis was blocked by the addition of 50 μg/ml kanamycin or 30 μg/ml chloramphenicol. Equal volumes of samples were collected at different time points as indicated in figures for Western analysis.

### Western analysis

Culture samples withdrawn at different time points were centrifuged. After removal of supernatant, the pellets were resuspended in 2X SDS sample buffer containing 40 mM DTT. The samples were boiled at 95 °C for 10 minutes and centrifuged to remove cellular debris. After centrifugation, the extracts were resolved on 10-12% SDS-PAGE gels. Proteins from the gel were then transferred to a polyvinylidene difluoride (PVDF) membrane. After blocking the membranes with 3% milk-TBST buffer, the membranes were probed with primary antibodies overnight at 4 °C. Antibodies used were: polyclonal rabbit anti-CtrA (1:5000 dilution), anti-DnaA (1:10000 dilution), anti-TacA (1:10000 dilution), affinity purified anti-PdeA (1:1000 dilution), anti-ClpP (1:5000 dilution) or monoclonal mouse anti-FLAG M2 (1:5000 dilution; Sigma). After washing off the excess primary antibody, the membranes were probed with goat anti-rabbit (Millipore, USA) or goat anti-mouse (Millipore, USA) secondary antibodies conjugated to HRP enzyme. Proteins were visualized by the luminescence from HRP substrate using a chemiluminescence detection system G-box (Syngene, UK).

## RESULTS

### Degradation of CtrA is enhanced in cells lacking the polar organizing protein PopZ

PopZ is a scaffolding protein that facilitates polar localization of a multitude of proteins including those that are directly involved in CtrA degradation such as CpdR, RcdA, PopA and ClpX (21, 29). Cells lacking PopZ have morphological defects and fail to localize the aforementioned factors to the stalked pole (21, 29). Since polar localization of the ClpXP protease and the substrate CtrA, facilitated by the cofactors CpdR/RcdA/PopA, was postulated to be critical for CtrA degradation, we hypothesized that CtrA degradation would be lost in cells lacking PopZ if this model was correct. Contrary to this hypothesis, we observed that CtrA was degraded even more rapidly in Δ*popZ* than in wildtype cells (Fig. 1A and 1B). CtrA stability and cell morphology were restored to those of wildtype cells when this Δ*popZ* strain was transformed with a plasmid expressing the protein PopZ, suggesting that the enhanced degradation of CtrA in Δ*popZ* strain is due to specifically to loss of PopZ (Fig. 1A, 1B and 1C). Together, these results suggest that the scaffolding protein PopZ restrains CtrA degradation and support our *in vitro* work where CpdR, RcdA, PopA-cdG physically interact even in the absence of subcellular localization to stimulate ClpXP-mediated degradation of CtrA (Fig. 1D and (4)).

**Figure 1.**
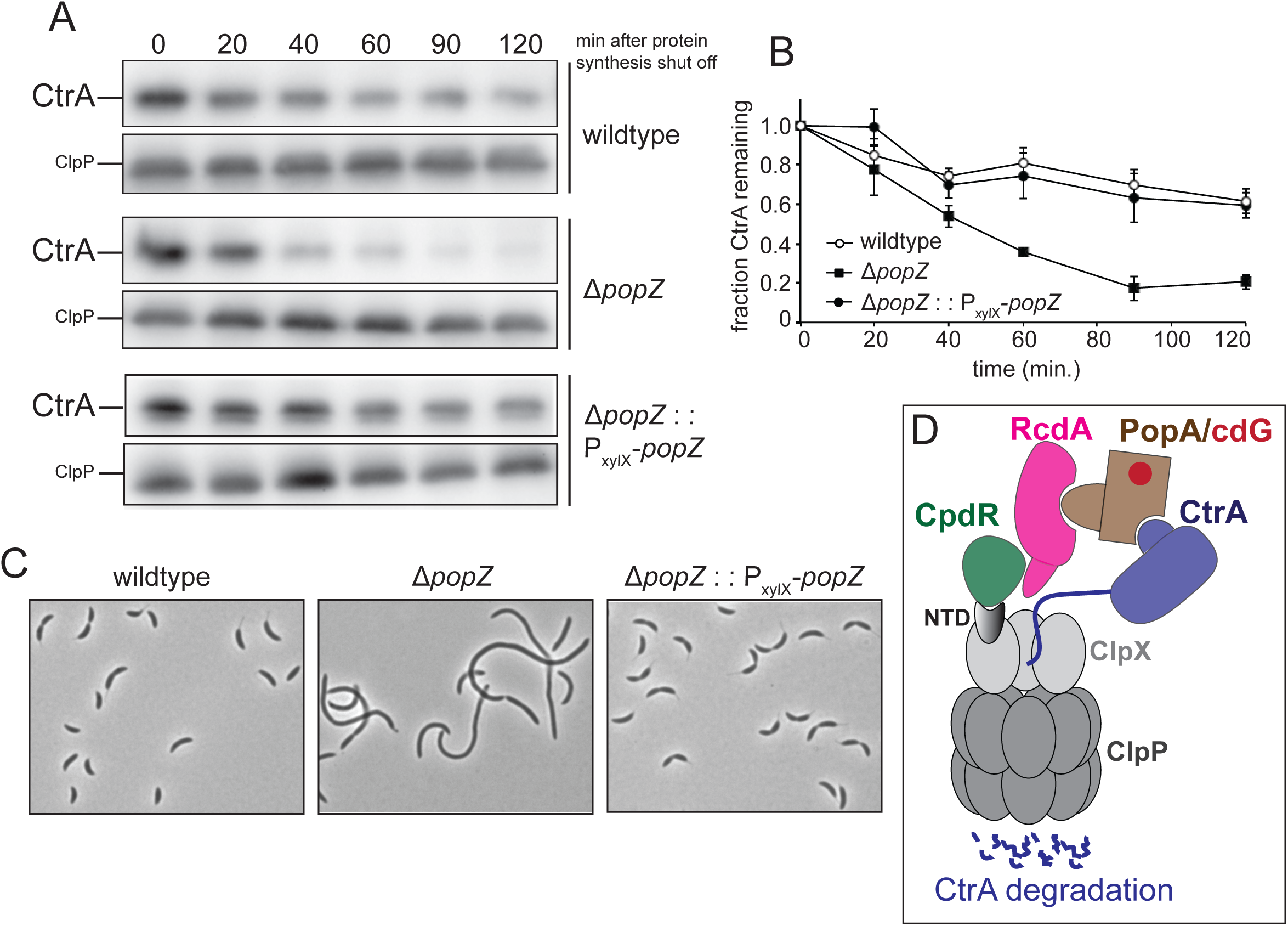
CtrA degradation is enhanced in cells lacking the pole organizing protein PopZ. (A) CtrA degradation in wildtype, Δ*popZ* and Δ*popZ* complemented by a PopZ expressing plasmid (n.b., the xylose promoter is leaky and PopZ is expressed even without addition of inducer). Cells were grown to exponential phase in PYE and then translation was blocked by adding kanamycin. CtrA stability was monitored by Western blot analyses by probing the blots with anti-CtrA antibody. ClpP is shown as a loading control. (B) Quantitation of Western blots. Bands corresponding to CtrA and ClpP were quantified using ImageJ (NIH, USA) and normalized band intensities over time are shown. Data represents mean ± SD of two independent experiments. (C) Complementation of PopZ protein expressed from a plasmid restored wildtype morphology of Δ*popZ* cells. All strains were grown to logarithmic phase in PYE. (D) A model depicting adaptor complex-mediated proteolysis of CtrA by ClpXP. Here CpdR, RcdA and PopA assemble in a hierarchical manner to deliver many substrates including CtrA to ClpXP for degradation (4).

### Adaptor-independent ClpXP and non-ClpXP proteolysis is not affected in cells lacking PopZ

The loss of PopZ protein stimulated CtrA degradation. This could be explained by prolific activation of ClpXP or proteolysis in general, or by activation of the adaptors needed to degrade CtrA *in vivo*. To address this, we monitored degradation of a ClpXP reporter substrate comprised of GFP fused to the C-terminal degron of CtrA (GFP~CtrA_15_) that does not require adaptors for degradation. Degradation of M2 epitope tagged GFP~CtrA_15_ is not affected in Δ*popZ* cells compared to wildtype cells suggesting that the stimulation of CtrA degradation is specific to changes in adaptor activity (Fig. 2A and 2B). To examine whether protein degradation is globally stimulated in Δ*popZ* cells, we monitored degradation of a ClpXP-independent substrate DnaA. Degradation of DnaA was unaffected in Δ*popZ* cells compared to wildtype cells suggesting that the enhancement in degradation in Δ*popZ* cells is specific to ClpXP-dependent substrates (Fig. 2C and 2D). Therefore, we next set out to examine if changes in adaptor activity explained the increased CtrA degration in Δ*popZ* cells.

**Figure 2.**
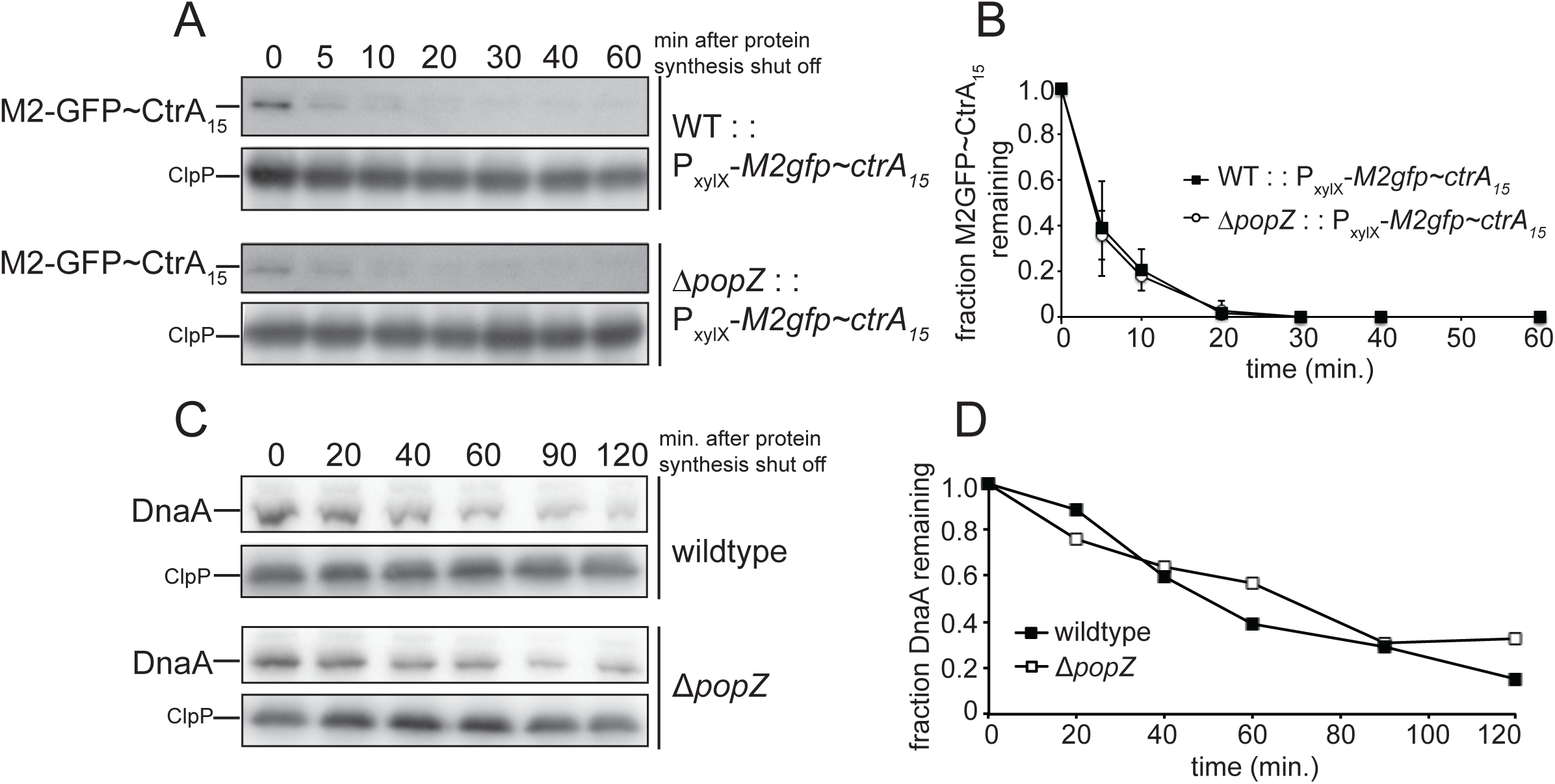
Degradation of either adaptor or ClpXP-independent proteolysis is not compromised in wildtype and Δ*popZ* cells. (A) Wildtype and Δ*popZ* cells expressing an M2-epitope tagged GFP~CtrA_15_ were grown to exponential phase and induced with 0.3% xylose for 2 hours. Protein synthesis was then blocked by the addition of chloramphenicol. Lysates from equal volumes of cells were collected at indicated time points for SDS-PAGE gels. M2-GFP~CtrA_15_ stability was monitored by Western blot analyses by probing the blots with a anti-M2 antibody. ClpP is shown as a loading control. (B) Quantitation of Western blots. Bands corresponding to M2GFP~CtrA_15_ and ClpP were quantified using ImageJ and normalized band intensities over time are shown. Data represents mean ± SD from two biological replicates. (C) Degradation of ClpXP-independent substrate DnaA is not affected in wildtype and Δ*popZ* cells. Conditions used were similar as in figure 1A except that the blot was probed for anti-DnaA antibody. ClpP is shown as a loading control. (D) Quantitation of Western blots from figure 2C.

### Degradation of both CpdR and RcdA-dependent substrates is enhanced in cells lacking PopZ

Because CtrA requires CpdR, RcdA and PopA for degradation, we explored the need for each tier of the adaptor hierarchy by using substrates specific to each level. PdeA is a phosphodiesterase that only requires the CpdR and TacA is a response regulator that requires both CpdR and RcdA for ClpXP-mediated degradation. (4, 15, 30). In order to determine which tier of adaptor-dependent proteolysis is compromised in Δ*popZ* cells, we monitored the degradation of both PdeA and TacA. Degradation of both PdeA and TacA is enhanced in cells lacking PopZ suggesting that along with CtrA degradation; degradation of other adaptor-dependent ClpXP substrates is also stimulated in Δ*popZ* cells (Fig. 3A and 3B). These results point to a conclusion that ClpXP is constitutively activated at the CpdR adaptor level.

**Figure 3.**
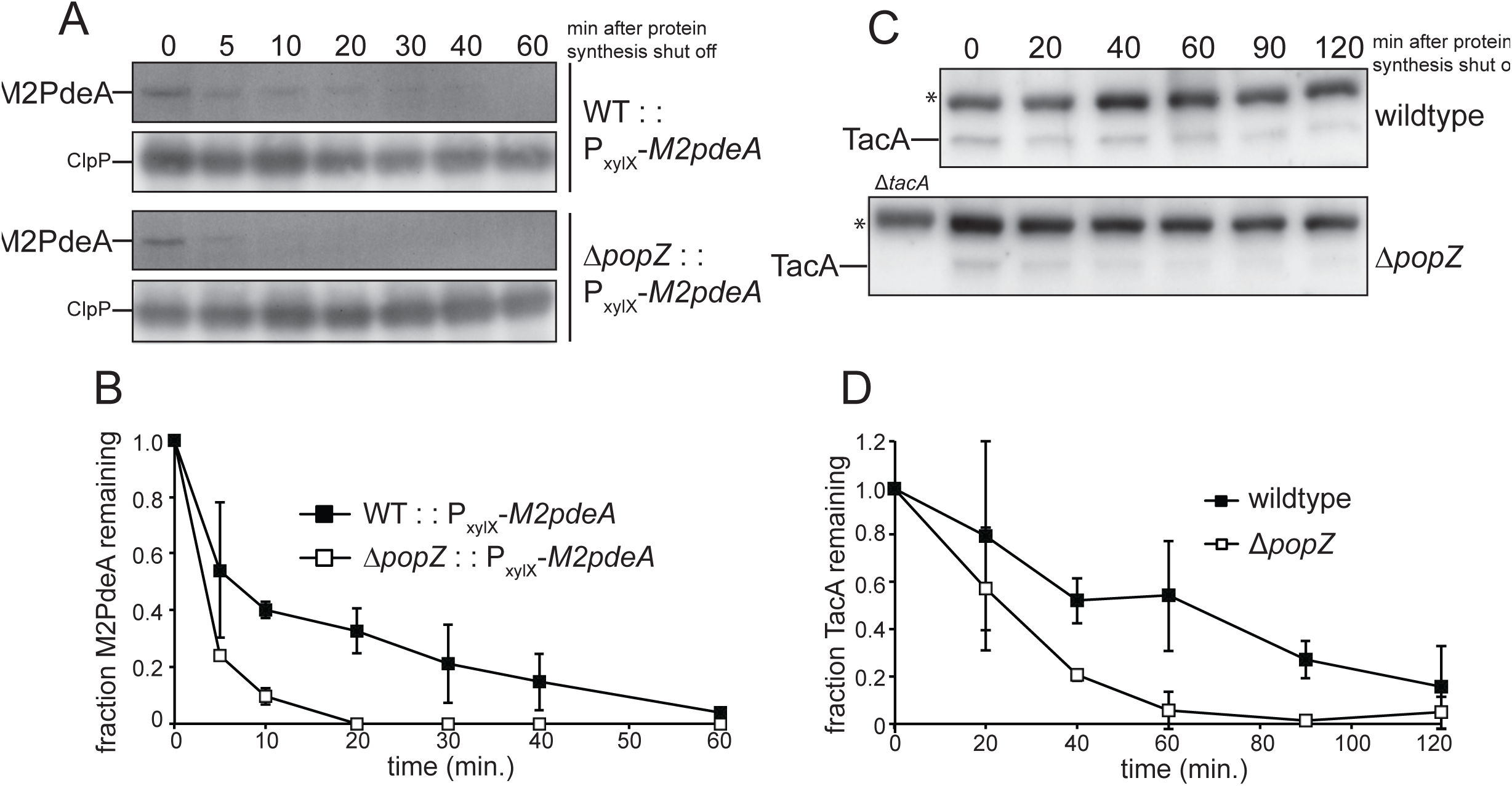
Degradation of CpdR and RcdA-dependent substrates is enhanced in cells lacking PopZ. (A) Strains expressing M2-tagged PdeA were grown to exponential phase and induced with 0.3% xylose for 2 hours before inhibiting protein translation by the addition of chloramphenicol. (C) TacA stability was monitored in wildtype and Δ*popZ* cells. Lysates from equal volume of cells were collected at indicated time points and loaded onto SDS-PAGE gels. PdeA and TacA stability was monitored by Western blot analyses by probing the blots with anti-M2 and anti-TacA antibodies. Asterisks denote cross-reacting band. (B, D) Quantitation of Western blots. Bands corresponding to M2PdeA, TacA and ClpP were quantified and normalized band intensities over time are shown. Data represents mean ± SD from two biological replicates.

### CpdR is epistatic to PopZ in CtrA degradation pathway

The adaptor CpdR is required for stimulation of CtrA degradation both *in vivo* and *in vitro* (4, 13, 14). We reasoned that since CpdR appears to be constitutively in Δ*popZ* cells, deletion of CpdR in this background should result in loss of CtrA degradation. To test this hypothesis we made a Δ*cpdR*Δ*popZ* double knock out strain by transducing Δ*cpdR:tet* into the Δ*popZ* strain using ϕCr30 phage. As expected, CtrA degradation was stabilized in Δ*cpdR*Δ*popZ* double knockout background similar to that in Δ*cpdR* strain (Fig. 4A). Microscopy experiments confirmed delocalization of CpdR-YFP in the Δ*popZ* and Δ*cpdR*Δ*popZ* cells (Fig. 4B and (21)). CtrA degradation was restored by complementing Δ*cpdR*Δ*popZ* strain with a plasmid expressing the CpdR protein suggesting that the loss of CtrA degradation in Δ*cpdR*Δ*popZ* strain could be attributed solely to loss of CpdR (Fig. 4C). CpdR activity is regulated by phosphorylation as a nonphosphorable CpdR (CpdR-D51A) prolifically delivers CtrA for degradation (13). Consistent with this, expression of CpdR-D51A in Δ*cpdR*Δ*popZ* strain increases CtrA degradation even more than WT (Fig. 4C). Taken together, these results suggest that the adaptor protein, CpdR, is absolutely required for CtrA degradation in Δ*popZ* cells.

**Figure 4.**
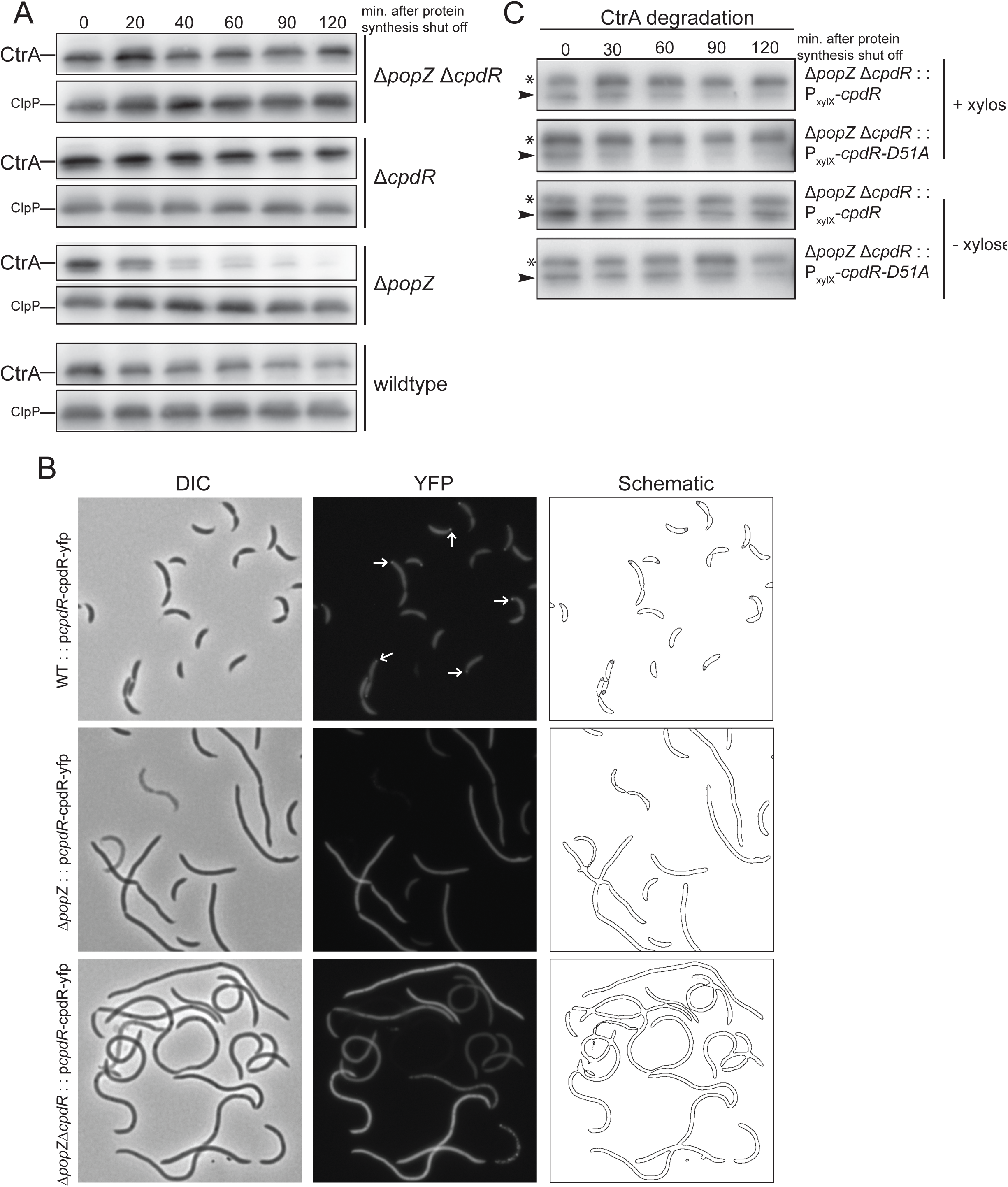
CpdR is epistatic to PopZ in CtrA degradation pathway. (A) CtrA stability is monitored in wildtype, Δ*popZ*, Δ*cpdR and* Δ*cpdR* Δ*popZ* strains. (B) Phase contrast and fluorescence microscopy images of wildtype, Δ*popZ*, Δ*cpdR* Δ*popZ* cells expressing CpdR-YFP from a plasmid bearing a CpdR promoter. Arrows indicate polar CpdR-YFP foci. (C) CtrA degradation is restored in Δ*cpdR*Δ*popZ* double knockout cells expressing CpdR from a plasmid. CpdR and CpdR-D51A were expressed in Δ*cpdR*Δ*popZ* cells from low copy xylose-inducible plasmids. Cells were grown to exponential phase with 0.3% xylose or not and protein synthesis was inhibited by adding chloramphenicol. Lysates from equal number of cells were collected at indicated time points for Western analysis. Asterisks denote cross-reacting bands.

### Overexpression of the PopZ protein stimulates CtrA degradation in a CpdR independent manner

Overexpression of PopZ leads to the enlargement of the polar region and over-recruitment of proteins such as CtrA, CpdR, RcdA and ClpX to these polar zones (21). To determine if prolific polar recruitment affects substrate degradation, we monitored CtrA degradation in strains overexpression PopZ. Because we had found that loss of PopZ results in increased CtrA degradation, we were surprised to find that overproduction of PopZ also results in faster CtrA compared to wildtype cells (Fig. 5A and 5B). This stimulation was specific to CtrA; PdeA degradation was not affected (Fig. 5A and S1). Overexpression of PopZ did not compromise protein degradation globally as degradation of a ClpXP-independent substrate, DnaA, remained unaffected (Fig. 5A and S1).

**Figure 5.**
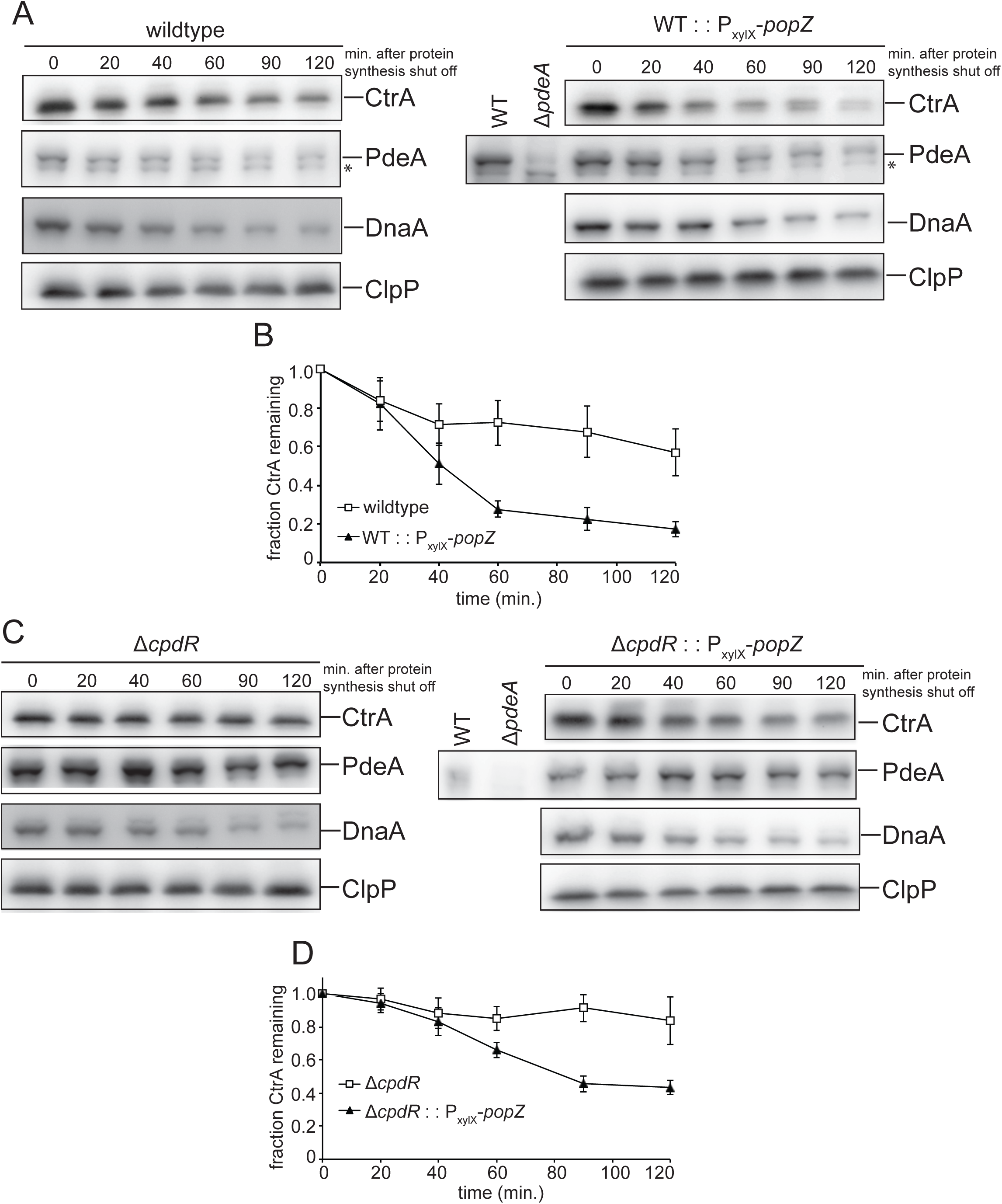
Overexpression of PopZ specifically stimulated CtrA degradation in wildtype and Δ*cpdR* cells. (A) Degradation of CtrA, PdeA, and DnaA was monitored in wildtype cells or WT cells overexpressing PopZ from a high copy xylose-inducible plasmid. Cultures were induced for 8 hours using 0.3% xylose keeping them in exponential phase all time. After inhibiting protein synthesis by the addition of kanamycin, lysates from equal volumes of cells were collected at indicated time points and loaded onto SDS-PAGE gels. Asterisks denote crossreacting bands. (B) Bands corresponding to CtrA and ClpP were quantified using ImageJ (NIH, USA) and normalized band intensities over time are shown. Data represents mean ± SD of three independent experiments. (C) Degradation of CtrA, PdeA, and DnaA was monitored in Δ*cpdR* cells or Δ*cpdR* overexpressing PopZ. (D) Bands corresponding to CtrA and ClpP were quantified and normalized intensities are shown. Data represents mean ± SD of three independent experiments.

Why is CtrA degradation stimulated upon PopZ overexpression but PdeA is not? We considered this result in light of the fact that PdeA absolutely requires CpdR for degradation both *in vivo* and *in vitro* (15, 31) while ClpXP alone can degrade purified CtrA *in vitro* (32). Our working model is that overexpression of PopZ forces ClpX and CtrA to the poles to increase local concentration sufficiently to allow for the recognition of CtrA directly by ClpXP. If this is true, then this overexpression should bypass the need for CpdR. Indeed, we found that CtrA, which is stable in Δ*cpdR* cells as expected, was degraded in Δ*cpdR* cells overproducing PopZ (Fig 5C and 5D). Importantly, PdeA was stable in the absence of CpdR, regardless of PopZ overexpression. Together these results suggest that forcing recruitment of the protease and the substrate by overexpressing PopZ is sufficient to bypass the need for adaptors when the substrate can be directly recognized by the protease.

## DISCUSSION

In this work, we explored if polar localization of the ClpXP protease is critical for its activation in *Caulobacter*. Prior genetic experiments showed that the adaptors CpdR/RcdA/PopA are necessary for normal CtrA degradation *in vivo* and microscopy-based experiments showed these factors facilitated localization of the ClpXP protease and the CtrA substrate to the stalked pole presumably to aid in degradation (13, 18, 19). Recent *in vitro* reconstitution experiments showed that all these accessory factors work together as biochemical adaptors enhancing the affinity of CtrA for ClpXP (4, 14). We found that CtrA is degraded rmore rapidly in cells lacking the PopZ protein compared to wildtype. The fact that the degradation of other CpdR and RcdA-dependent substrates is also enhanced in Δ*popZ* cells supports a model where polar localization mediated by PopZ protein might be critical for restraining adaptor-mediated protein degradation rather than driving degradation.

CckA is a membrane-bound bifunctional histidine kinase. Autophosphorylation of CckA results in phosphorylation of both CtrA and CpdR via the phosphotransferase ChpT (10–12). When bound to cdG, CckA switches to a phosphatase preferring state and dephosphorylates CtrA and CpdR (16, 17). This allows cyclic fluctuation of CtrA activity via phosphorylation and proteolysis, which is key for proper cell cycle progression. Because increased local density of CckA at the stalked pole appears to drive its kinase activity (33) and PopZ is needed for CckA localization (23), a working model for our results are that loss of CckA localization forces it into a phosphatase state resulting in constitutive dephosphorylation of CpdR and resulting activation of adaptor dependent ClpXP degradation of substrates. Consistent with this, cells expressing mutant CckA alleles with disrupted polar localization result in more rapid CtrA degradation (34).

Our results combined with others support a model where polar localization organized by PopZ normally influences degradation of ClpXP substrates through regulating the activity of CckA rather than by forcing substrates in close proximity with proteases (Fig. 6). However, artificially increasing PopZ can force corecruitment of ClpXP with substrates to drive degradation even in the absence of adaptors. When PopZ is present at normal levels, CckA can switch between the kinase and the phosphatase states to tightly control CpdR-dependent ClpXP activation. However, when PopZ is deleted, CckA predominantly functions as a phosphatase, resulting in dephosphoryation of CpdR and constitutively activation of adaptor dependent ClpXP proteolysis. Interestingly, recent work shows that certain stresses lead to rapid clearance of CtrA through a switch in CckA to its phosphatase state without changes in CckA localization (35), further supporting the critical role in CckA activity rather than its localization per se in controlling CtrA degradation.

**Figure 6.**
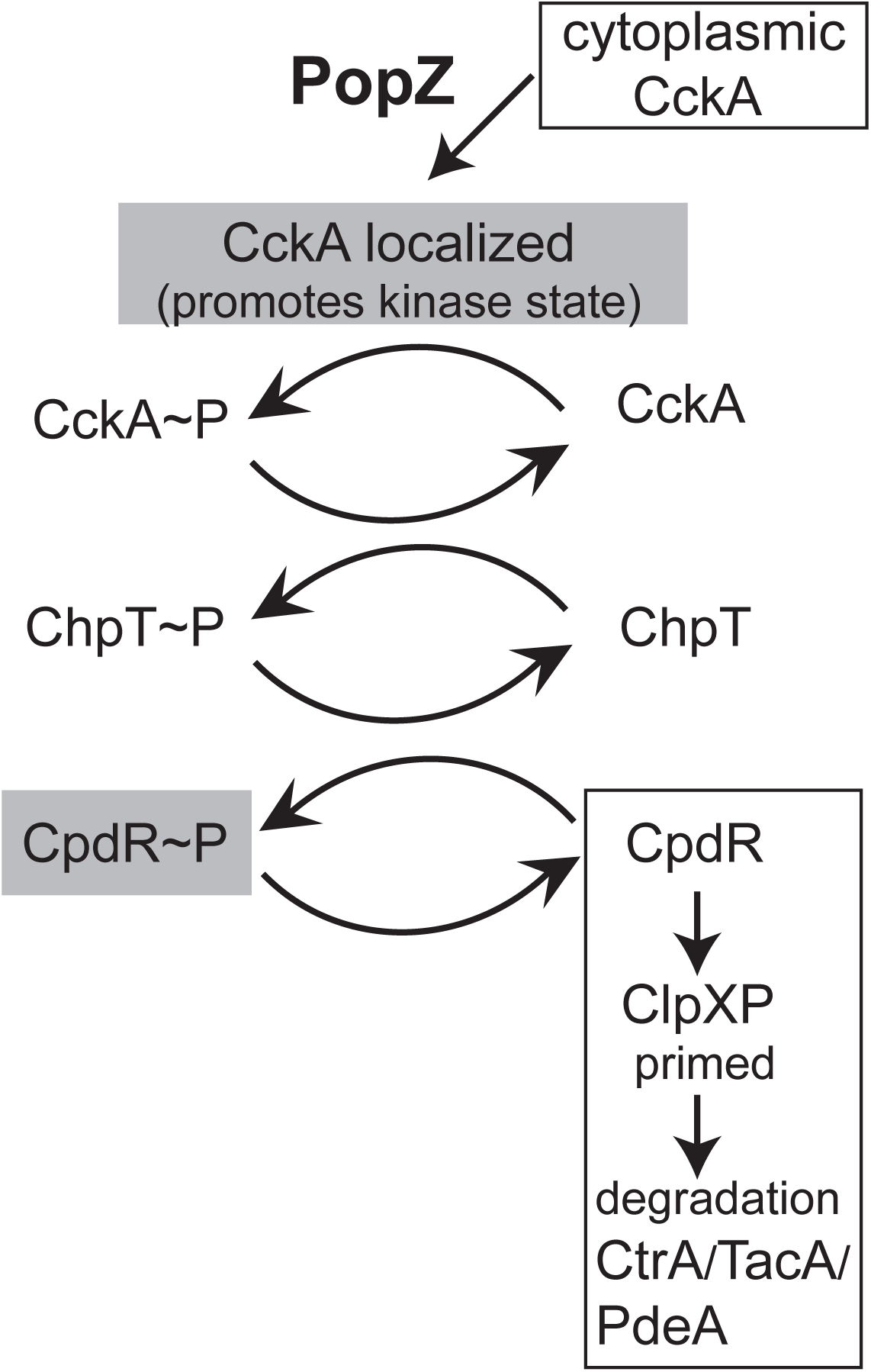
A model representing the role of PopZ is restraining proteolysis. When PopZ is present, CckA is properly localized to stalked pole where it can switch between a kinase and a phosphatase states thus modulating phosphorylation of CpdR. When CpdR is dephosphorylated it can prime the ClpXP protease to activate degradation of substrates CtrA/TacA/PdeA. When PopZ is absent, CckA is delocalized and predominantly functions as a phosphatase (17, 34). CckA then dephosphorylates CpdR via ChpT. The dephosphorylated CpdR constitutively primed ClpXP protease and prolifically stimulates degradation of CtrA/TacA/PdeA.

## ACKNOWLEDGEMENTS

We thank Chien lab members for helpful discussions. K.K.J thanks Amrita Palaria for helpful comments and discussions. We also thank Christine Jacobs-Wagner for providing PopZ expression plasmids and strains and Patrick Viollier for graciously providing anti-TacA antibody. Work in the Chien lab is supported by NIH grant R01GM111706 (to P.C).

## Supplemental information

**Figure S1.**
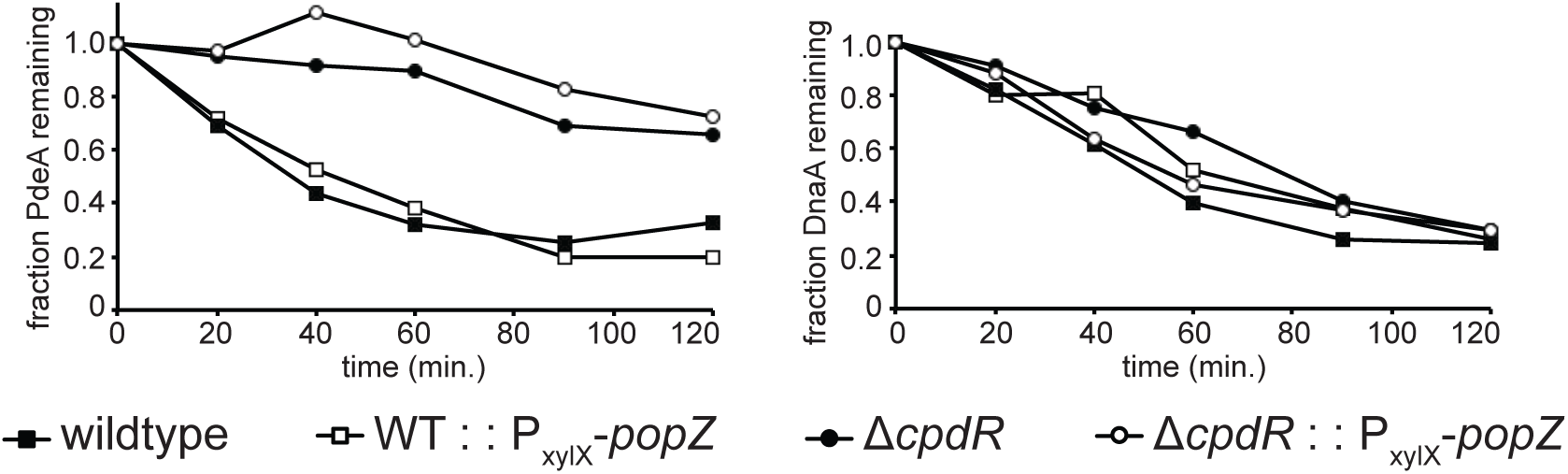
Quantitation of Western blots from Figure 5. Bands corresponding to PdeA and DnaA were quantified using ImageJ and normalized band intensities over time are shown.

